# Efficient and accurate prediction of protein structure using RoseTTAFold2

**DOI:** 10.1101/2023.05.24.542179

**Authors:** Minkyung Baek, Ivan Anishchenko, Ian R. Humphreys, Qian Cong, David Baker, Frank DiMaio

## Abstract

AlphaFold2 and RoseTTAFold predict protein structures with very high accuracy despite substantial architecture differences. We sought to develop an improved method combining features of both. The resulting method, RoseTTAFold2, extends the original three-track architecture of RoseTTAFold over the full network, incorporating the concepts of Frame-aligned point error, recycling during training, and the use of a distillation set from AlphaFold2. We also took from AlphaFold2 the idea of structurally coherent attention in updating pair features, but using a more computationally efficient structure-biased attention as opposed to triangle attention. The resulting model has the accuracy of AlphaFold2 on monomers, and AlphaFold2-multimer on complexes, with better computational scaling for large proteins and complexes. This excellent performance is achieved without hallmark features of AlphaFold2, invariant point attention and triangle attention, indicating that these are not essential for high accuracy prediction. Almost all recent work on protein structure prediction has re-used the basic AlphaFold2 architecture; our results show that excellent performance can be achieved with a broader class of models, opening the door for further exploration.

## Introduction

RoseTTAFold (RF) ^1^ was developed before the architecture of AlphaFold2 (AF2) ^2^ had been made public. While both models were dramatic increases over state-of-the-art protein structure prediction at release, AF2 had considerably higher accuracy overall. RF and AF2 differ in both their architecture and training regimen. In terms of architecture, major differences between RoseTTAFold and AF2 consist of: a) the inclusion of a third “3D structure” track in the RF basic block, b) the use of biaxial attention in the 2D pair track (RF) versus triangle multiplication and triangle attention (AF2), c) the use of the SE3-equivariant transformer ^3^, in RF rather than Invariant Point Attention (IPA) in AF2 for computing structural updates, and d) the overall depth of the network, with 8+4 2-track/3-track layers in RF versus 4+48 full-msa/clustered-msa evoformer layers in AF2. In terms of training, AF2 added: a) the idea of “recycling,” where inputs from one network pass were fed into the next, with a random number of recycles used as “preconditioning” on each training example, b) the use of frame-aligned point error (FAPE) as the structure loss function, and c) the use of distillation data, where highly confident network outputs were fed in as new training examples. While several of these changes (recycling and distillation) were explored in ablation studies ^2^, it is unclear which features are critical for the high accuracy of AF2, and whether an alternate architecture could achieve comparable accuracy.

With the full release of the AF2 method, we set out to combine the best features of both models, and to determine what features were required for the remarkable AF2 prediction accuracy. We reasoned that by adding in features from AF2 one by one, and assessing whether prediction accuracy increased, we could distinguish those that were essential to high accuracy prediction. Since the original publication of RF and AF2, multiple methods for accurate protein structure modeling have been described that essentially re-use the exact AF2 architecture with some modification of inputs: OpenFold ^4^ is a reimplementation of AF2; ESMFold ^5^ replaces the multiple sequence alignment (MSA) with a language model generating a single featurized sequence; OmegaFold ^6^ develops a new pair-to-pair update as a simplified version of triangle attention but makes use of IPA; and UniFold ^7^ uses AF2 architecture with modifications to loss and auxiliary heads. We reasoned that building a new model from scratch would provide more insight into the critical determinants of accurate structure prediction and guide future efforts in this field; we also aimed for improvements in efficiency and to develop a single model with state of the performance on both monomer and assembly prediction. The resultant model, RoseTTAFold2 (RF2) is a single model equivalent in accuracy to AF2 for monomers and AF2-multimer for complexes, with better computational scaling on proteins and complexes larger than 1000 residues. Finally, our RF2 model – along with inference and training code – is made freely available.

## Results

An overview of the RF2 network architecture is shown in Figure 1. Like the original RoseTTAFold, we maintain three parallel tracks, representing the 1D sequence/MSA, 2D residue-pair, and 3D protein structure features. These blocks are repeated 36 times; following AF, the first four blocks make use of a larger set of 1024 sequences as MSA inputs, while the remaining blocks use only 128 sequences. The training procedure was largely unchanged from RF with a few exceptions: a) 0-3 iterations of the network were run as a preconditioning step, with network weight updates computed only from the final iteration; b) in addition to monomers, the network was trained on all contacting pairs of chains from biological assemblies; and c) a distillation set of 1.2 million high-confidence AF2 predictions was added to the training set. We evaluate the model by comparing on a large set of recent Protein Data Bank (PDB) depositions (released following model training) of monomers (113 medium and hard CAMEO ^8^ targets released between 2023-01-14 and 2023-04-08) and complexes (140 hetero-dimers released after 2021-09-30), and compare performance to AF2 and AF2-multimer (release v2.3.1) in Figure 2a-b. Network running time as a function of protein length (on a single A100 GPU) is shown in Figure 2c. The modeling performance on CASP14 targets was tracked as features were added, removed and modified (Figure 2d). The remainder of this section describes each of the modifications made from RF in building RF2, and the effect of these on modeling performance for CASP14 targets, focusing on novel model features and the improvements that led to the largest performance gains.

**Figure 1.**
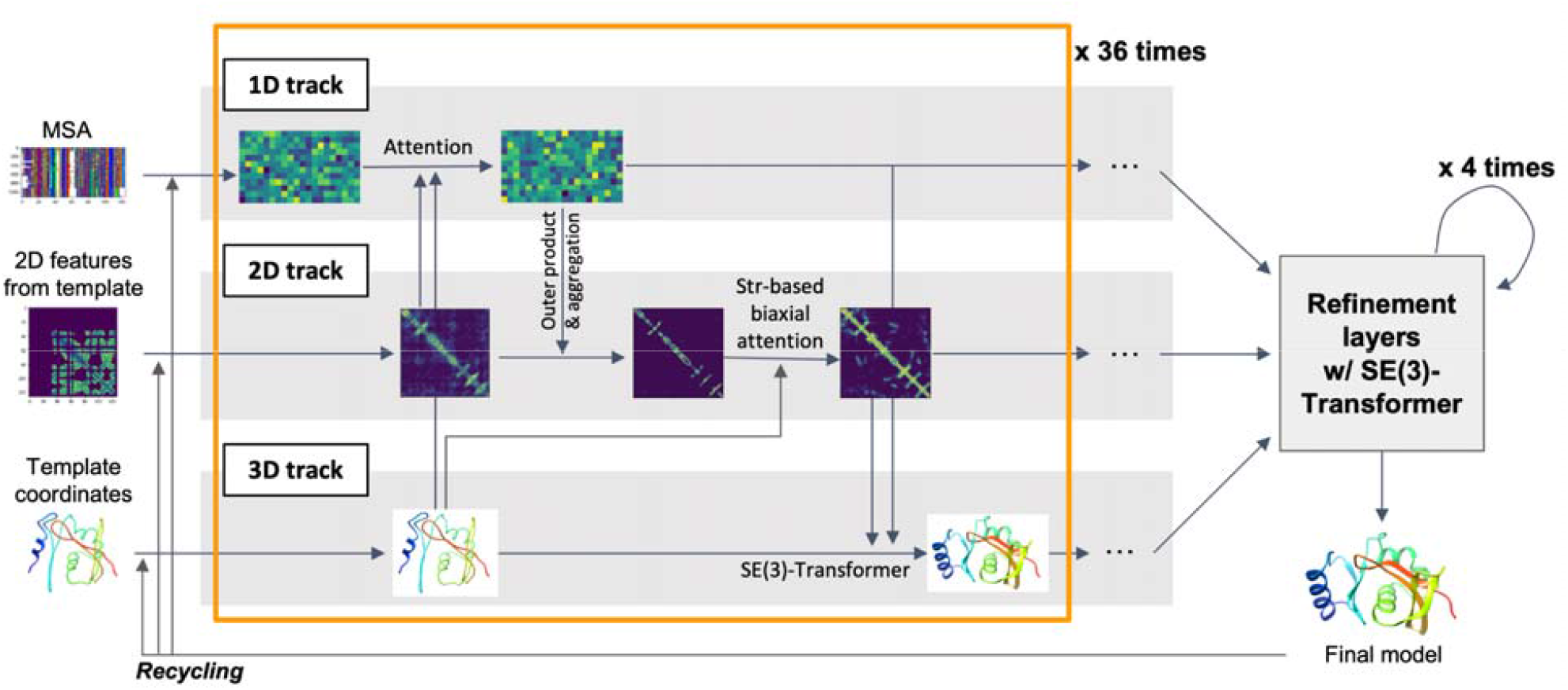
An overview of the RF2 3-track architecture. Three parallel tracks synchronize and update a representation of the sequence, residue-pair, and 3D structure, producing a fine 3-dimensional structure.

**Figure 2.**
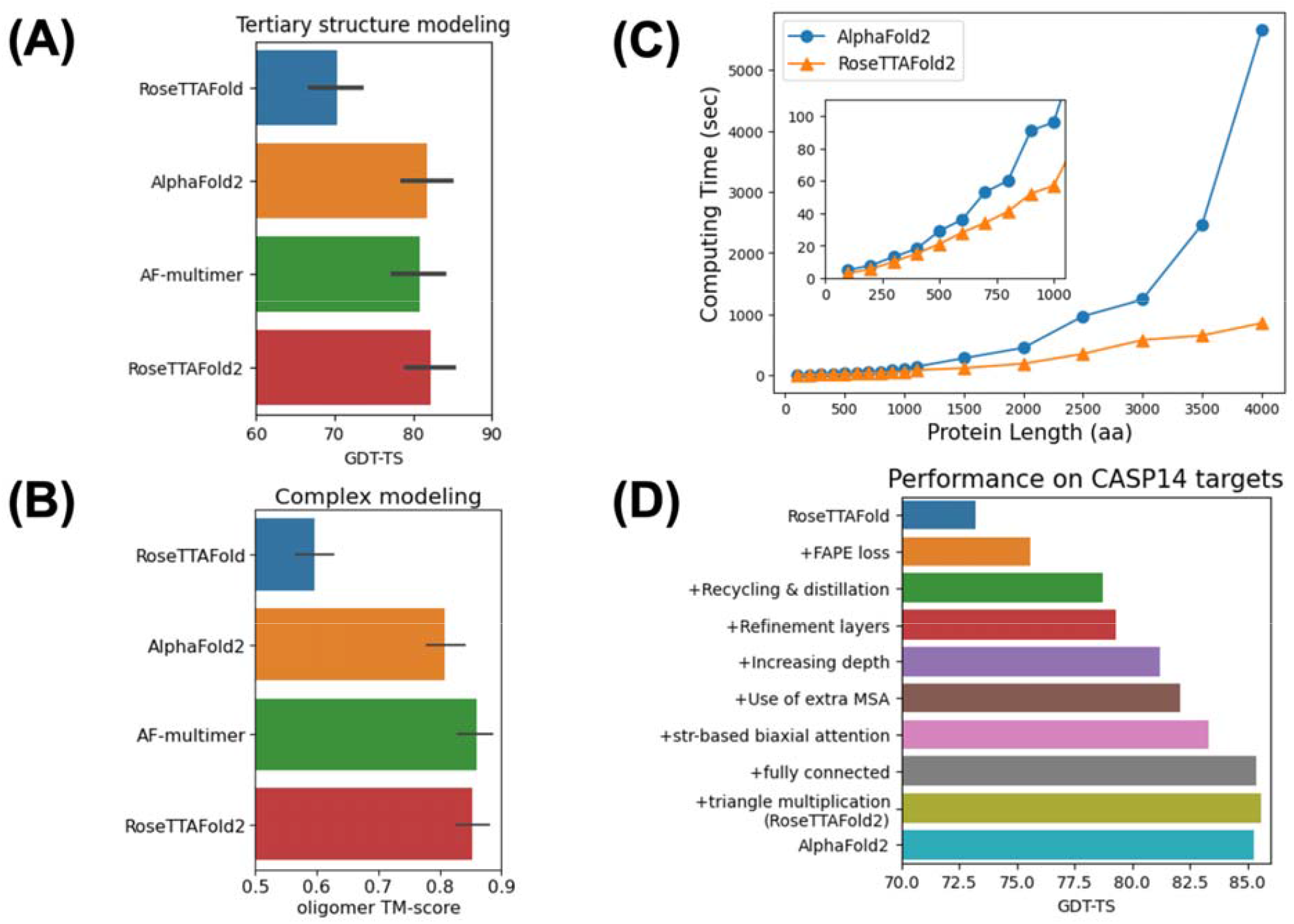
Comparison of the results of RF2 versus AF2. On a set of recently-solved structures (released after training of AF and RF2), we compare the average accuracy of structure prediction over A) 113 protein monomers and B) 140 protein hetero-dimer complexes. C) A comparison of AF2 and RF2 in terms of runtime scaling. D) CASP14 performance as individual features were added to the model.

AF2 utilized triangle attention in the pair to pair updates to implicitly bias pair features towards residue-pair distances respecting 3D locality, without needing to tightly tie predictions to predicted distance histogram (“distogram”) losses. However, triangle attention is computationally expensive for large proteins, requiring attention calculations over all *ijk* residue triplets. With RF’s 3D track providing structure information at every level of the network, we hypothesized that this 3D structure track could provide a more efficient route to incorporating 3D locality information into the 2D pair track. We experimented with adding a structure-derived bias to the biaxial attention of RF2, using residue-pair distances from the current 3D structure to bias attention calculations in the pair-to-pair updates ^9^. This could allow the network to encode the same inductive bias as triangle attention, but without the computational expense of triangle attention. Following extension of the 3D structure track along the full length of the model, we found that using residue-pair distances derived from the 3D track to bias the attention weights in the pair-to-pair updates led to clear model improvements: a 48-layer model with the old RF architecture (36 2-track + 8 3-track + 4 shared refinement layers) was outperformed by a 3-track-only model (12 3-track + 4 shared refinement layers) with structure-biased attention and one quarter the total parameters. Increasing the depth of 3-track layers from 12 to 36 further improved the performance, improving the average GDT-TS over CASP14 targets by 2.0.

AF2 introduced an invariant-point attention (IPA) module to generate 3D coordinates from MSA and pair features; in RF, this role was carried out by the SE(3)-equivariant transformer. We found that when used in the three-track architecture of RF, IPA led to unstable training performance. In RF2, we use the same SE(3)-equivariant transformer as the structure generation component, but made slight modifications. Following AF2, we introduced an additional SE(3)-Transformer-based “refinement” layer which takes the MSA, pair embeddings, and backbone coordinates from the final 3-track block and generates updated backbone and sidechain coordinates over four iterations. This led to a GDT-TS improvement over CASP14 targets by 0.6.

At this point, our model had similar average accuracy to AF2, but was worse for large, 300+ residue, “easy modeling cases;” that is, protein families with relatively deep MSAs. In some of these cases, the main deviations were in the relative positions of regions separated by long distances, resulting from very subtle “domain bending” relative to the native structure. We hypothesized this could be due to structure updates only passing information to the 128 nearest residues. Allowing direct information flow between all residues (and adjusting the number of layers and channels so memory use was similar), resulted in improved performance for such targets; the average GDT-TS increased 2.2 when structure updates were fully connected. This fully-connected geometry was only used in the main 3D track of the model, with the refinement block limited to passing information to the 64 nearest residues.

We obtained further improvements with incorporation of several additional features of AF2. The use of FAPE loss in place of the average of distance- and coordinate-RMS in RF led to a 2.4 unit improvement in GDT-TS over CASP14 targets. Recycling – running multiple iterations of the network for each prediction – and using AF2 predictions ^10^ to augment the training set, improved the GDT-TS by 3.1. Incorporating more network layers, with fewer channels per layer also proved beneficial: increasing from 12 to 48 layers while keeping the number of parameters the same led to a 1.9 GDT-TS improvement. Updating the input embeddings to use up to 2048 sequences (rather than the 256-sequence limit in RF); and including triangle multiplication in addition to biaxial attention in the pair-to-pair updates improved the CASP14 GDT-TS by 1.9 and 0.2, respectively.

## Discussion

Our development of RF2 from RF by incorporating features from AF2 highlights the features of AF2 that contribute most to its high accuracy. We find that while certain features from AF2 were critical (at least in our experiments) for achieving high accuracy – including FAPE loss, recycling, distillation, and network depth – other features, like IPA and Triangle attention, can be replaced by alternate architectures without reducing accuracy. FAPE loss in particular was very important in giving smooth gradients towards the native structure from the entire conformational space, which superimposed Cα RMS and distance-map RMS both do not – the former suffers from numeric stability issues when the prediction is far from the target, and the latter has an alternate minimum at the mirror image. IPA on the other hand is one of many equally-good solutions to the problem of updating a rotationally invariant point-cloud guided by 1- and 2-body features. Many recent efforts have made use of AF2’s evoformer and IPA architecture in carrying out various protein structure prediction and design tasks ^11–14^; RF2 provides an orthogonal structure prediction framework to AF that enables *in silico* screening of designs.

While the performance of RF2 is not more accurate than AF2 in our tests, it should be useful to have a single model with state-of-the-art performance on both protein monomer and complex modeling, and with better scaling to large systems. For challenging structure modeling problems, having multiple distinct high accuracy methods to build models should also be useful. RF2 is freely available at https://github.com/uw-ipd/RoseTTAFold2.

We have found that RF can be readily extended to protein design by fine-tuning for inpainting ^15^ and diffusion tasks ^16^. Having a structure track that extends to the input enables prior information (for example, a target structure to design binders to, or an enzyme active site geometry) to be encoded as “multi-body” structures rather than pairwise distances and orientations (e.g., ^15^). Such direct encoding in 3D coordinates is very natural for design constraints, and we speculate that this could contribute to better performance in design challenges, but we have not systematically explored pure two-track models without such a 3D structure track.

## Methods

### Network architecture

The network architecture of RF2 – illustrated in Figure 1 – is similar to that of RF ^1^. The network consists of three “tracks”: a 1D MSA track, a 2D residue-pair track, and a 3D structure track. Each track is initialized through a series of embedding layers from the initial MSA and template features, and subsequently modified by embedding the recycled MSA and pair features. The recycled 3D structure is directly passed into the 3D track. Then, 36 rounds of the main “iteration” are carried out, where the 1D, 2D, and 3D tracks are sequentially updated from their previous values and the values of the other tracks. Finally, four iterations of the 3D update are carried out using frozen 1D and 2D data from the last layer of the network. Using a similar mechanism to AF2, we “recycle” the outputs from the previous network pass, using the embedding network described above. Below, we describe individual components of the network in detail.

#### Feature embedding

The initial features for the 1D, 2D, and 3D tracks – *msa, pair*, and *xyz*/*state*, respectively – are initialized from the input msa (*msa*_*in*_) and template structure (*xyz*_*in*_), as well as the previous iterations’ features (*msa*_*prev*_, *pair*_*prev*_, *xyz*_*prev*_, and *state*_*prev*_) in three steps. Step 1 computes the initial *msa, pair, xyz*, and *state* features from the inputs:

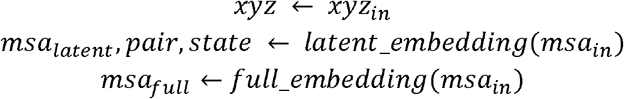

Step 2 modifies this information from previous iterations:

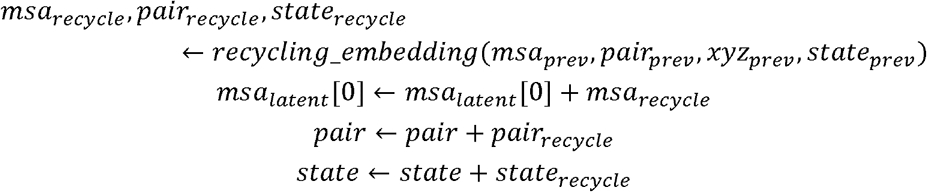

Finally, step 3 updates the 2D and 3D tracks with information from up to four template structures, with sequence *seq*_*template*_ and coordinates *xyz*_*template*_.

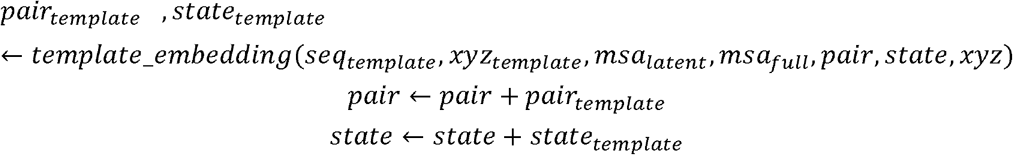

The *latent_embedding* module consists of the following:

- MSA features are embedded using linear layers mapping (N x L x 22) dimensional inputs to (N x L x d_msa_) outputs. An addition (L x 22) to (L x d_msa_) mapping is carried out on the input sequence and is added to every sequence in the mapping.
- Pair features are comprised of a “left” and “right” embedding of the input sequence from (L x 22) to (L x d_pair_); an outer sum of these builds the full (L x L x d_pair_) pair tensor, to which a positional embedding is added. Following AF2, the positional encoding bins *sequence* distance from -32 to +32, mapped to dimension d_pair_ via a linear layer.
- State features (for the 3D track) embed the (L x 22) input sequence to (L x d_state_) embedded dimension.

RF2 uses 256, 128, and 32, for d_msa_,d_pair_, and d_state_, respectively, and the number of sequences *N* is limited to 256. The *full_embedding* module is similar with two key differences: there is no positional embedding, and the full MSA (up to 2048 sequences) is used instead.

The *recycling_embedding* module uses the previous network iterations’ 1D, 2D, and 3D features to update the corresponding tracks in the current iteration:

- MSA and state features are normalized and returned (where they are added to the current values)
- Pair features are updated from the previous pair features as well as residue pair distances from the current model *xyz*_*prev*_. An (L x L x 64) tensor contains – for each residue pair *i,j*, the binned residue-pair distance from 0-32Å (0.5Å bin width) concatenated to the state features of *i* and *j*. This is embedded to dimension *d*_*pair*_, and added to the normalized pair features.

Finally, the *template*_*embedding* is the most complicated among the embedding procedures. In RF2, this involves converting *M* input templates into 1D and 2D features: the 1D features, *template*_*1D*_ contain the template sequence (1-hot encoded) concatenated with a per-residue confidence (as predicted by HHpred) and the sine and cosine of (up to four) template sidechain torsions, while the 2D features, *template*_*2D*_, contain residue pair distance and orientation following trRosetta ^17^. Specifically, a binned distance (Cβ-Cβ) is concatenated with the cosine and sine of 3 angles (Cα-Cβ-Cβ, and the dihedrals N-Cα-Cβ-Cβ and Cα-Cβ-Cβ-Cα) describing residue pair orientation. These features, *template*_*1D*_, and *template*_*2D*_ are then embedded and added to *pair* and *state* through the following procedure:

- *template*_*1D*_ is projected to *d*_*template*_ channels, and attention is applied across template structures, with *state* as the query and *template*_*1D*_ as the key/value pair.
- *template*_*2D*_ is left- and right-concatenated to *template*_*1D*_ and projected to *d*_*template*_ channels. This tensor is then subject to 2 iterations of biaxial attention along the sequence (identical to the pair-to-pair updates below) followed by attention across template structures (as with *template*_*1D*_)

These features are returned and added to *state* and *pair*.

#### Main iteration block

The main iteration block of the network is analogous to AF2’s “Evoformer,” and is used to update the 1D, 2D, and 3D tracks of the network using the previous values in the same track as well as information from the other tracks. This block is repeated 40 times, though – following AF2 – the first four iterations use the outputs of the *full_embedding* module, while the remaining 36 use the *latent_embedding* model outputs. The main update block consists of the following updates to features at step *i* (*msa*_*i*_, *pair*_*i*_, *xyz*_*i*_, and *state*_*i*_) from those at step i-1:

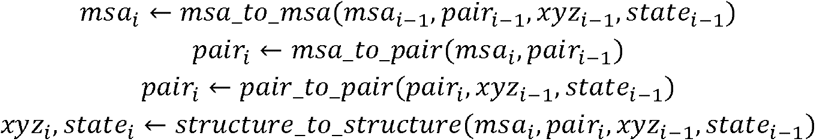

All the updates are quite similar to those used in the original RF, so they are only briefly highlighted below, with the differences versus the original implementation highlighted.

The *msa_to_msa* update uses the same biaxial attention as in the original RF implementation, where attention is first taken over residues, and then over sequences. However, two modifications have been made. First, state features are used to modify the query sequence (*msa*[0]) by projecting from (L x d_state_) to (L x d_msa_) and adding to *msa*[0]. Second, *pair* and *xyz* features are used to bias the attention across residues:

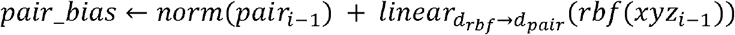

Here, *rbf* computes an (L x L x d_rbf_) matrix with the binned distances between residues *i* and *j* (convoluted by a Gaussian of width 1 grid point). When computing attention across residues, this pair bias is added to the attention calculation:

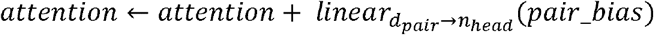

The *msa_to_pair* update is identical to that of the original RF, where the msa features are twice projected to d_hidden_ (16 in RF2), the outer product of both projections are taken, and this result is projected to d_pair_ and added to *pair*.

The *pair_to_pair* update is also similar to the original RF implementation (which used the architecture of the MSATransformer ^9^) but with two key additions. First, the structure from the 3D track is used to bias both attention calculations (row and column) in the pair update, though the functional form is a bit different (0 denotes outer product):

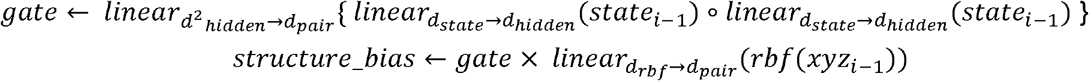

As above, *structure_bias* is used as a bias on both attention calculations. The second change from the original implementation is the use of AF2’s triangle multiplication prior to the biaxial attention calculations, where “in” and “out” multiplications were carried out with a hidden dimension of 32.

Finally, the *structure_to_structure* update is related to the original RF implementation, where an SE(3)-equivariant transformer ^3^ is used to update a set of coordinate frames representing the protein backbone (N-Cα-C). Unlike the original implementation, there is no initial graph transformer to initialize the coordinates, and the SE(3) transformer is used on the input structure. The input structure comes from a template – if one is provided – with all unaligned residues placed randomly within 5Å of the closest unaligned residue. If no input template is provided, all residues are randomly placed within 5Å of the origin. Unlike the original implementation, the graph used in SE(3) is fully connected; attention and message updates are computed over all residues.

Additionally, sidechain prediction is added to the network. The RF2 sidechain prediction module is nearly identical to the AF2 implementation with several modifications. First, in working with the SE(3)-equivariant transformer rather than AF2’s IPA, instead of having a set of state features used to generate both backbone updates and sidechain updates, the SE(3)-equivariant transformer calculates two vector (*l*=1) features, giving the backbone updates, and 32 scalar (*l*=0) features, from which sidechain conformations are computed. The ResNet architecture and angle encoding as sine/cosine pairs of AF2 was used, with additional prediction of two “bending” angles around the Cα atom around the Cβ angle, allowing some “non-ideality” in the protein structure, and leading to a total of 10 predicted angles per residue. This sidechain prediction is applied at every level of the network, but the predictions are not directly “fed forward,” only indirectly via the state features.

#### Refinement Block

After 40 iterations of the main block, four more rounds of the *structure_to_structure* are applied. These four repeats use shared parameters, and – compared to the normal structure-to-structure updates – increase the number of channels and state features (channels increased to 128 from 48; state features increased to 64 from 32), and only compute attention from each residue to its 64 nearest neighbors. These changes are intended to allow the network to adjust fine details of the structure as opposed to making large structural changes.

#### Auxiliary Heads

In addition to predicting structure from MSA, the network also predicts several auxiliary features from one or more of the three tracks:

- Amino-acid logits are predicted from the final layer of the 1D (MSA) track, projecting d_msa_ to 21
- Distograms and orientations are predicted from the 2D (pair) track, computing binned predictions for the distance and three angle terms described above. This auxiliary head projects d_pair_ to d_feat_ for each subterm (d_feat_ is 19 for the angle term, 37 for the others) and symmetrizes the two torsional terms so θ_ij_=θ_ji_.
- pAE (following AF2) – the estimated error per residue pair – is computed from the pair features, by projecting d_pair_ to 64 pAE bins (of width 0.5)
- pLDDT – the estimated lddt per residue – is computed from the state features, projecting d_state_ down to 50 lddt bins
- pBind – the probability that two chains are binding – is computed from pAE logits. It averages pAE over all interchain cells and projects down from the 64 bins to a single value and takes the sigmoid to produce a single pBind value (per interface)

### Network Training

#### Training dataset

The dataset used for training consists of all protein structures in the PDB published before April 30, 2020. Compared to the original RoseTTAFold, our processing pipeline was a bit more permissive, including all crystallographic and cryo-electron microscopy structures solved to a resolution of 4.5Å or better, as well as all NMR structures. Following RF, structures were split into individual chains, and clustered using an e-value cutoff of 1e-3, yielding 280k structures in 20k clusters.

A “distillation” set of protein monomers was also created using AF2 for structure prediction. 12M sequences in UniRef50 were folded by AF2 ^10^, and a subset of high-confidence (>0.8 pLDDT) and long (>200 residues) predictions were selected. We cluster as before, yielding 3.6M sequence/structure pairs in 1.0M clusters. Protein chains sharing >30% sequence identity to any of the protein chains from the PDB-derived training set were excluded.

Finally, we created a dataset of multimers for training. Using the biological assembly annotations, for each chain in our training dataset, we identify and annotate all chains in contact with that chain in a biological assembly, separately annotating homomeric and heteromeric interactions. This is then divided into two prediction tasks during training: for *heteromeric* training, we provide the network pairs of contacting chains, and have the model recover their specific dimeric conformation; for *homomeric* training, we provide two copies of the chain, and have the model recover any dimeric conformation present in the biological assembly (e.g, for a D2-symmetric homomer the model is correct in recovering *either* two-fold interface, with backpropagation pulling toward the closer of the two).

#### MSA and template generation

MSA generation for all targets is identical to that of RF: HHblits ^18^ is run against UniRef30 with e-value cutoffs of 1e-10, 1e-6, and 1e-3, respectively, stopping when either 2000 sequences with >75% coverage or 4000 sequences with >50% coverage are found. If these criteria have not been met even at an e-value of 1e-3, a search is then run against BFD ^19^. Templates are generated by running HHsearch ^20^ with up to 500 templates with HHsearch probability > 5%. Templates were not generated for the distillation data (both in initial structure generation and in retraining); only MSA information was used.

#### Loss function and training parameters

The loss function in RF2 draws heavily from that used in training AF2, though with several key differences. A listing of individual loss terms and their weights are highlighted below:

- **wt**_**disto**_=1.0, the distogram loss, takes the cross-entropy between predicted distograms and the native
- **wt**_**aa**_=3.0, the masked AA prediction loss, takes cross entropy between the actual sequence and the predicted sequence over masked positions only (see below for masking procedure)
- **w**_**str**_=10.0, the structure loss, split equally between: a) average backbone FAPE loss over the predictions at all 44 structure updates, and b) average all-atom FAPE of the final predicted structure.
- **w**_**pae**_=1.0, the loss on the networks’ pAE prediction, taken as cross-entropy between predicted and true pAE
- **w**_**plddt**_=1.0, the loss on the networks’ pLDDT prediction, taken as cross-entropy between predicted and true pLDDT
- **w**_**bond**_=0.02, the loss on bond geometry. The sum of losses on peptide bond deviation from the ideal, consisting of a length term (C-N) and two bond-angle terms (Cα-N-C and N-C-Cα). These loss terms are flat in a narrow range (±Δ) around an ideal value, and are quadratic outside of this window. For the length term, an ideal length of 1.329 and a Δ of 0.02 is used; the angle terms are calculated in cosine space, where ideal values of - 0.4415 and -0.5255 are used, with a Δ of 0.05. Finally, these terms are normalized by the number of violations made.
- **w**_**lj**_=0.02, the clash loss. This term uses Rosetta’s ^21^ Lennard-Jones (LJ, 12-6) potential, normalized by the number of residues. This term provides a weak attraction between atoms as they approach their ideal distance, and a strong repulsive force as they get too close. In RF2, we use a slightly weaker version of Rosetta’s potential, setting *lj_lin* (a parameter controlling when the potential goes linear) to 0.75 compared to its default of 0.6.
- **w**_**bind**_=1.0, the predicted binding loss. The binary cross-entropy between the pBind head and whether a pair of chains comprises a complex.

Training is run using a 50%/25%/25% split between distillation data, PDB monomers/homoligomers, and PDB heterooligomers. When a homoligomeric PDB is encountered in training, it has a 50% chance to be modeled as a complex. Within the heterooligomer dataset, 50% of the time non-binding protein pairs are provided; in these cases, structure loss is computed over chains individually (and pBind is driven to 0). Following AF2, MSAs are randomly masked, with 15% of all sequences chosen for masking, with 70% given a specific mask token and the remaining 30% changed to a different amino acid. Training with recycling also follows AF2, with 0-3 random passes through the network run on each example, before a final iteration through which backpropagation is enabled. Large proteins/protein complexes are randomly cropped. For monomers, a continuous stretch of residues is carried out, while, for heterooligomers, a “structure-aware” cropping is carried out that prefers interface residues.

Training is run in two phases: in the first, bond and LJ loss weights are set to 0, and crop size is set to 256. The Adam optimizer is used, with a learning rate of 1e-3, and L2 weight decay. Approximately 100,000 minimization steps (with a batch size of 64) are carried out. In the second stage, bond and LJ weights are set to 0.02, crop size is set to 384, and the learning rate is reduced to 5e-4. An additional 50,000 minimization steps are carried out.

## Acknowledgments

We thank Microsoft for their generous donation of Azure compute time, and Meta for pre-publication access to 12 million AF2 predictions.

This work was supported by the New Faculty Startup Fund from Seoul National University.

